# Temporal dissociation of salience and prediction error responses to appetitive and aversive taste

**DOI:** 10.1101/184341

**Authors:** E. Hird, W. El-Deredy, A. Jones, D. Talmi

## Abstract

The feedback-related negativity, a frontocentral event-related potential (ERP) occurring 200-350 milliseconds (ms) after emotionally-valued outcomes, has been posited as the neural correlate of reward prediction error, a key component of associative learning. Recent evidence challenged this interpretation and has led to the suggestion that this ERP expresses salience, instead. Here we distinguish between utility prediction error and salience by delivering or withholding hedonistically matched appetitive and aversive tastes, and measure ERPs to cues signalling each taste. We observed a typical FRN (computed as the loss-minus-gain difference wave) to appetitive taste, but a reverse-FRN to aversive taste. When tested axiomatically, frontocentral ERPs showed a salience response across tastes, with a particularly early response to outcome delivery, supporting recent propositions of a fast, unsigned and unspecific response to salient stimuli. ERPs also expressed aversive prediction error peaking at 285ms, which conformed to the logic of an axiomatic model of prediction error. With stimuli that most resemble those used in animal models we did not detect any frontocentral ERP signal for utility prediction error, in contrast with dominant views of the functional role of the feedback-related negativity ERP. We link the animal and human literature and present a challenge for current perspectives on associative learning research using ERPs.

## Introduction

The reward prediction error hypothesis of associative learning provides a foundational understanding of adaptive behaviour and is used widely to explain the neuroimaging correlates of associative learning in humans (Holroyd & Coles, 2002; O’Doherty, Hampton, & Kim, 2007). The seminal finding that sparked much of this research is that midbrain dopamine neurons express reward prediction error, increasing their firing to unexpected reward and reducing their firing to unexpected omission of reward (Schultz et al., 1997), with an impressive homogeneity in the firing of individual dopamine neurons (Eshel, Tian, Bukwich, & Uchida, 2016). The term ‘reward’ is inherently related to the term ‘utility’, an economic term that denotes the subjective value of an outcome, from ‘good’ to ‘bad’ (Friedman & Savage, 1952). Reward prediction error specifically refers to the signal the brain is thought to compute when it encounters an unexpected delivery or omission of an appetitive outcome, but the literature often assumes that reward prediction error signals actually reflects the utility of either appetitive or aversive outcomes (Schultz, 2016). This assumption means that unexpected delivery of appetitive outcomes should be signalled similarly to the unexpected omission of aversive outcomes.

This interpretation of the midbrain dopamine signal has been challenged by evidence that some dopamine neurons respond with phasic bursts to the delivery of *both* appetitive and aversive outcomes, suggesting that they code salience, instead of utility (Brischoux, Chakraborty, Brierley, & Ungless, 2009; Bromberg-Martin, Matsumoto, & Hikosaka, 2010; Joshua, Adler, Mitelman, Vaadia, & Bergman, 2008). Schultz (2016) proposed that the challenge salience presents to the interpretation of midbrain dopamine firing as a reward prediction error can be addressed by distinguishing between two temporally separate signals within 500 ms of a predictive cue: an initial nondiscriminative response to various forms of salience, between 100-200ms from the cue, followed by a utility prediction error signal from 150-350ms (Schultz, 2016). Schultz (2016) listed three reasons for this initial salience response. Physical salience refers to physical attributes such as size and colour, motivational salience refers to the ability of an outcome to elicit attention due to its high motivational relevance, while surprise salience refers to the unexpectedness or novelty of an outcome. Clearly, both appetitive and aversive outcomes can have high motivational and surprise salience (Sambrook & Goslin, 2014). Higher salience of stimuli increases their ability to capture attention. For example, an unpleasant taste is low in utility and so could elicit a negative reward prediction error, but is also high in motivational salience, because it represents a potentially harmful substance which would attract attention so that it is avoided in the future (Esber & Haselgrove, 2011). A negative reward prediction error would be expressed by a reduction in neural activity, whereas a salience response would be expressed by an increase.

Distinguishing between utility and salience is fundamental in the burgeoning literature in neuroeconomics and affective neuroscience. This is evident in recent meta-analyses of human neuroimaging (Bartra, McGuire, & Kable, 2013; Lindquist, Satpute, Wager, Weber, & Barrett, 2015; Sambrook & Goslin, 2015a). The doubt regarding what dopamine neurons compute has triggered a reexamination of the functional role of a key event-related potential (ERP) that is thought to be dopaminergically mediated. The feedback-related negativity (FRN) is thought to originate from dopaminergic projections to the anterior cingulate cortex (ACC), evident by the finding that it is modulated by dopamine agonists (Holroyd & Coles, 2002; Santesso et al., 2009; Walsh & Anderson, 2012) and combined EEG-fMRI work (Hauser et al., 2014). The dominant theory contends that the feedback-related negativity (FRN) expresses utility prediction error, but recent studies provided evidence that it expresses salience (Garofalo, Maier, & di Pellegrino, 2014; Hauser et al., 2014; Huang & Yu, 2014; Pfabigan et al., 2015; Sambrook & Goslin, 2015b; Talmi, Atkinson, & El-Deredy, 2013). Those studies showed that the FRN reflects a negative deflection when any outcome – appetitive or aversive – is unexpectedly omitted, as proposed by the predicted response-outcome (PRO) model (Alexander & Brown, 2011) and in line with an interpretation of the FRN as expressing motivational salience rather than utility prediction error signal.

A criticism of those studies is that most appetitive and aversive outcomes were not well-matched. For example, in two studies that used money and physical pain as reinforcers (Heydari & Holroyd, 2016; Talmi et al., 2013) the response latency of the frontocentral EEG signal differed (Heydari & Holroyd, 2016). While disparate spatiotemporal dynamics are unsurprising when appetitive and aversive outcomes have different hedonic values and are drawn from different modalities, as in both of these studies, such spatiotemporal differences make it difficult to argue that a signal expresses the psychological variable of salience in both appetitive and aversive conditions. Importantly, in traditional FRN studies that use financial outcomes, ostensibly matching modality and outcome value, monetary gain may represent a different level of motivational salience than monetary loss, so that the dimension of value could be confounded by that of salience (e.g. Bai, Katahira, & Ohira, 2015; Hauser et al., 2014; Weismueller & Bellebaum, 2016). Studies that employ financial outcomes rarely match the utility of reinforcement (e.g. Bai et al., 2015; Hauser et al., 2014) so that an FRN could stem from greater emotional and ensuing cognitive impact of losses over gains, as described in prospect theory (Sambrook, Roser, & Goslin, 2012). Indeed, there is evidence that midbrain neurons signal to negative more than positive prediction errors, suggesting negative prediction errors have greater salience (Rodriguez, Aron, & Poldrack, 2006). Because electrophysiological recordings demonstrate that different temporal ranges of the signal express different psychological variables, it is important to match outcome modality and utility.

A functional magnetic resonance imaging (fMRI) study that addressed the issue of salience and utility with well-matched outcomes, using taste reinforcers, observed a salience but not a utility prediction error response (Metereau & Dreher, 2013), in accordance with other fMRI studies that examined this distinction with other outcomes (Gu et al., 2016; Hauser et al., 2014). However, the recent realisation that differences between the dopaminergic signature of utility prediction error and salience can be discerned in different temporal ranges of the signal (Schultz, 2016) means that unique utility responses in those studies were perhaps lost to temporal smearing in fMRI work. Here we used EEG, which has greater temporal resolution, to test whether any ERP could be detected which expresses utility prediction error rather than salience across well-matched appetitive and aversive outcomes.

We used taste reinforcers, which closely resemble reinforcers used in the animal models that the associative learning literature uses as its key interpretative framework. Indeed, while previous research has demonstrated the feasibility of using taste in appetitive conditioning in EEG (Franken, Huijding, Nijs, & van Strien, 2011), ours is the first ERP study of taste prediction error. Importantly, we used hedonically equidistant sweet and bitter tastes, which circumvents the confound of utility and motivational salience present in traditional FRN studies and allows a direct comparison of the signal in the appetitive and aversive conditions, to distinguish between utility and salience response. We observed an FRN for appetitive outcomes, but a reverse FRN for aversive outcomes. Further analysis exploited the logic of the axiomatic model of prediction error (described under ‘Experimental Design’ below) to assess whether signal codes salience or utility prediction error (Roy et al., 2014; Rutledge, Dean, Caplin, & Glimcher, 2010). We found ERPs that expressed salience and an aversive prediction error; we did not observe an ERP expression of utility prediction error. These results are important because they contradict the dominant theory of the FRN and reveal the limits in what we can observe on the human scalp.

## Method

### Experimental design

Axiomatically, a neurobiological signal of **utility prediction error** should be expressed as a specific form of interaction between outcome and expectancy (Rutledge et al., 2010). When good outcomes (here, delivered sweet and omitted bitter taste) and bad outcomes (omitted sweet and delivered bitter taste) are expected, the signal should not differentiate between them, so expected outcomes form a baseline for comparison, in agreement with direct recordings of dopaminergic neurons (Schultz et al., 1997). The difference between good and bad outcomes should be pronounced when outcomes are unexpected. This logic of the original model requires a manipulation of valence and expectancy but not of outcome domain (appetitive/aversive). It would be enough, for example, to cross expectancy with delivered and omitted sweet taste, or with delivered sweet and delivered bitter taste. A manipulation of valence and expectancy is not enough, however, to convincingly show that a signal expresses utility prediction error. This is because a utility prediction error signal must express both appetitive and aversive prediction errors, namely, it must exhibit the interaction described above separately in the appetitive (here, sweet) and aversive (here, bitter) domain. Moreover, although it is not known whether *more* dopamine corresponds to *more* or *less* positive amplitude in any particular instance, if the funnel plot for the appetitive domain shows that unexpectedly delivered positive outcomes are more positive than unexpectedly omitted positive outcomes, the opposite should then hold in the aversive domain, where unexpectedly delivered negative outcomes should be *less* positive than unexpectedly omitted negative outcomes. Therefore, to investigate whether the ERP signal expresses utility prediction error or salience we manipulated taste: (bitter / sweet), expectancy (expected / unexpected) and outcome (whether taste is delivered / omitted) within subjects (Figure 1). This factorial design is appropriate to investigate whether any neurobiological signal conforms to the axiomatic model (Rutledge et al., 2010). A difference wave approach cannot tell us, for example, whether unexpected good outcomes diverge away from Expected outcomes in the opposite direction than unexpected bad outcomes.

**Figure 1:**
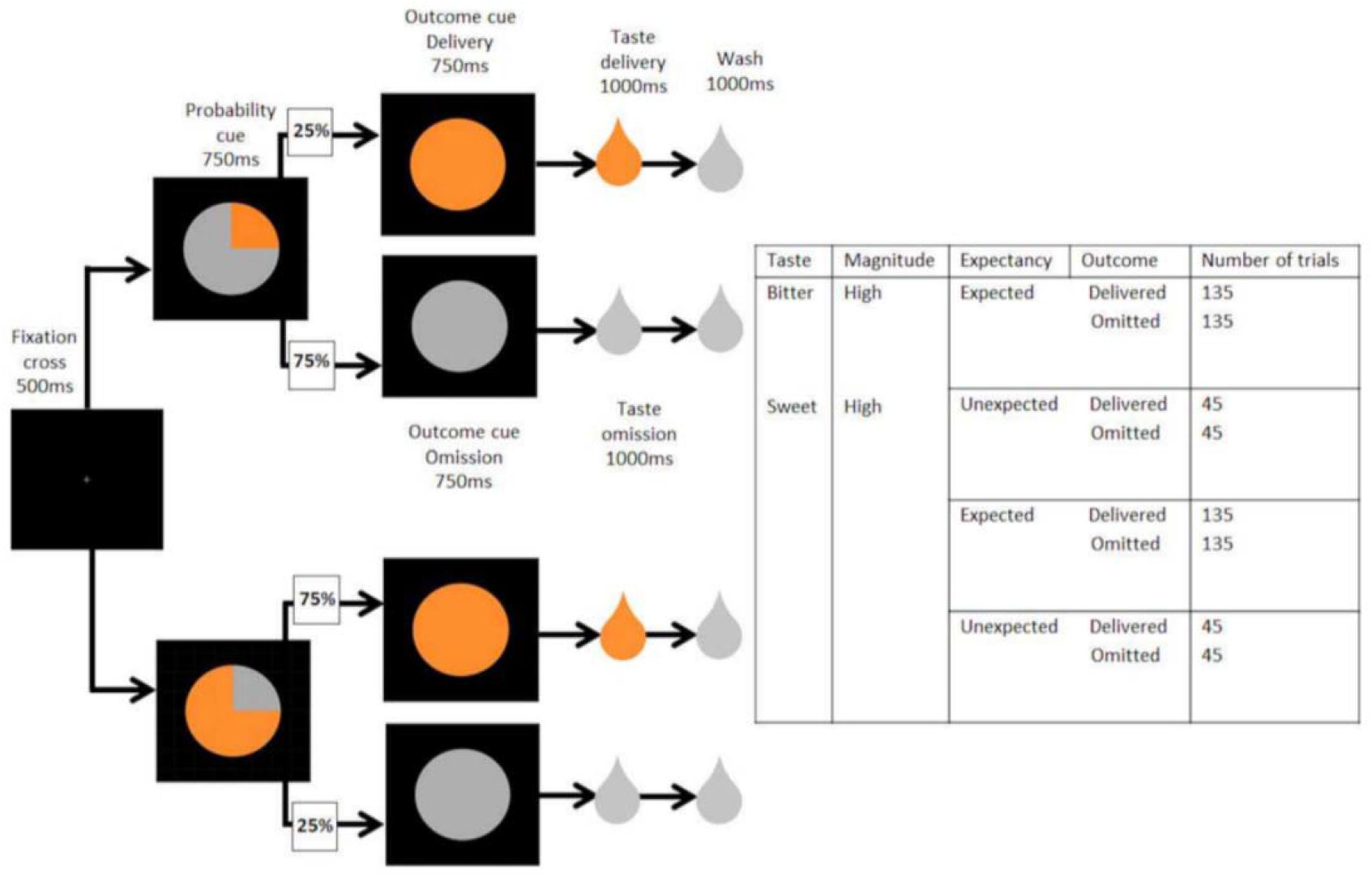
Experimental paradigm. Top: The timeline of each trial. After a fixation cross, a veridical probability cue set the expectation of the taste delivery/omission chance of the trial. An outcome cue followed, declaring the impending true outcome of the trial, deliver/omit, which triggered a theoretical prediction error. The taste was delivered, or omitted, based on the outcome cue. A rinse solution of artificial saliva prepared the participant for the next trial. *Bottom: Number of trials in each condition.* ‘Expected’ outcomes were presented 75% of the time, and ‘unexpected’ outcomes 25% of the time, to match the veridical probability cue.

**Motivational salience** is operationalised as a main effect of outcome, differentiating cues that predict delivered taste from those that predict omitted taste. **Surprise salience** is operationalised as a main effect of expectancy, differentiating cues that are expected from those that are unexpected. These main effects could be expressed regardless of whether the taste is appetitive or aversive. To maintain the perceived salience of experimental taste delivery, a subset of additional trials in the experiment delivered low magnitude tastes (slightly sweet, slightly bitter), which were administered in keeping with the same experimental design, but not analysed. Bitter and sweet tastes were delivered in separate blocks, to prevent reward generalisation (Schultz, 2016), namely a response to the bitter taste because participants were expecting the sweet taste.

### Participants

Twenty participants aged 18-35 (12 females, mean age 20 years) received £20 compensation or course credit points for participation in the study. Participants had normal or corrected-to-normal vision. They had no history of neurological or psychiatric conditions, no metabolic disorder and were not taking any centrally-acting medication or any medication which could make it difficult to fast for 2 hours. Ethical approval was granted by the University of Manchester, where the study took place.

### Materials

Pilot studies identified four concentrations of sucrose and water (sweet) and four concentrations of quinine and water (bitter), which elicited a range of hedonic ratings of pleasantness and unpleasantness, respectively. For sweet those were 1.0, 0.6, 0.2 and 0.1 mole of sucrose, and for bitter those were 1.0, 0.75, 0.25 and 0.05 millimole of quinine. Each solution also included 10 millilitres of lemon juice per litre of liquid to balance the sweetness of the sucrose whilst maintaining symmetry between sweet and bitter taste.

For each participant, two concentrations from each taste were selected, corresponding to equivalent high and low taste magnitude pairs, whose hedonic ratings were within 10% of one another (see procedure). The *low magnitude* hedonic ratings were at least 20% less than the *high magnitude* concentrations for that participant.

A neutral solution was designed to mimic the ionic balance of saliva to minimise reward associated with the liquid, and comprised of distilled water, 0.012 moles sodium bicarbonate and 0.012 moles potassium chloride.

Taste magnitude and likelihood were communicated to participants by visual cues, represented in figure 1, which were matched for luminance. The experiment was conducted on a Matlab platform (Mathworks).

### Apparatus

Participants’ heads were stabilized using a chin rest. Four rubber tubes [bore 2mm, wall 0.5mm, Altec Product LTD] were attached to the chin rest and to the participant’s face with medical tape. The tubes were attached to 50ml syringes (Plastipak syringe 50ml Luerlok, Fisher Scientific) fitted into pumps (Harvard Apparatus, pump 33,) in the neighbouring room. Pump activity was controlled by software (Matlab, Mathworks).

### Procedure

Participants made two visits to the lab, and were instructed to avoid food and water for two hours prior to each visit.

On the first visit, in order to establish equivalent behavioural responses in bitter and sweet, participants rated four concentrations of quinine and four concentrations of sucrose using the labelled magnitude scale (LMS), a validated scale for collection of intensity ratings (Hayes, Allen, & Bennett, 2013). The LMS is a quasi-logarithmically spaced verbally labelled line describing hedonic intensity from *strongest imaginable* to *no sensation*. Participants were instructed to place a mark on the line where their perceived intensity of pleasantness for sweet, or unpleasantness for bitter, lay. Bitter and sweet were administered in separate blocks. Block order was randomised. The taste stimulus was delivered for 1000 ms. Participants then rated the taste on the LMS and had a sip of water before moving on to the next taste. Participants rated each stimulus five times in each block. Trial order was randomised. On the basis of those ratings, four concentrations were selected for each participant (see Materials).

On the second visit, participants were sat in a quiet, dimly lit room. A total of 720 trials were administered in the high magnitude condition. Each trial lasted 10 seconds, and the entire session lasted 3 hours. Bitter and sweet were administered in separate, alternating and counterbalanced blocks. The different magnitude and probability conditions were presented pseudorandomly within each block.

Participants were advised at the beginning of a block whether it was a ‘sweet’ block, or a ‘bitter’ block. Figure 1 depicts a schematic of the design and a timeline of each trial. A fixation cross was presented for 500ms followed by a veridical probability cue presented for 750ms which set up expectations: either 75% probability of taste delivery and 25% taste omission, or 75% probability of taste omission and 25% taste delivery. Next, an outcome cue was presented for 750ms. The outcome cue signalled the actual outcome (with 100% probability) of either delivery (orange) or omission (grey) of the taste. The probability of the deliver/omit outcome cues followed the statistics of the expectation cues: On trials with 25% probability of taste delivery, the taste delivery outcome cue was presented 25% of the time, and on trials with 75% probability of taste delivery, the taste delivery outcome cue was presented 75% of the time. Next, the taste was delivered, or omitted, based on the outcome cue. On ‘omit’ trials a neutral solution was delivered. Both taste solutions and neutral solutions were administered for 1000ms. Finally, a wash of neutral solution for 1250ms prepared the participant for the next trial. There was a pause of 2500ms during which the screen was black before the next trial began (see figure 1).

Breaks were provided every 40 trials. Every 80 trials a randomly selected cue was presented to participants and they were asked to identify whether they had seen this in the last 40 trials. This was designed as a check that participants were paying attention to the images on the screen.

### EEG recording

Continuous EEG recording was acquired at a sampling rate of 512 Hertz using a 64 electrode Active-Two amplifier system (Biosemi, Amsterdam, Netherlands) with Actiview acquisition software (Biosemi, Netherlands). Here, an active and passive electrode replaces the ground electrode to create a feedback loop that drives the average potential of the subject (the common mode voltage) as close as possible to the analogue-to-digital converter reference voltage in the analogue-to-digital box. Vertical and horizontal electro-oculograms (EOG) was measured for detection of eye-movement and blink artefacts. Impedances were kept at 20 KΩ or less. The experiment was conducted in a dimmed, quiet room.

### EEG data analyses

#### Preprocessing

The ERP time-locked to the outcome cue was preprocessed using SPM12 (Ashburner et al., 2013; Litvak et al., 2011). The signal was re-referenced to the mean of all scalp electrodes, downsampled to 200 Hertz (Hz), and filtered with a Butterworth filter between 0.1 and 30 Hz. Epochs were extracted 200ms before the outcome cue to 600ms after, importantly avoiding the actual delivery of any fluid, which occurred 750ms after the outcome cue. Artefact rejection was achieved by following two steps. Firstly, eyeblinks were modelled and underwent artefact rejection at a lenient threshold of 150uV. The resulting eyeblinks model was used to correct for eyeblinks, using the singular value decomposition (SVD) technique implemented in SPM12. Any remaining trials in which the signal in any of the electrodes exceeded 80 μV were rejected. On average, 17% of trials were removed across participants and conditions. One participant was removed from the analysis due to high noise levels in the ERP signal. Single-trial data were averaged separately for the eight conditions using the “robust averaging” method in SPM12b (Litvak et al., 2010). Robust averaging takes into account distribution of data for every channel and trial by down-weighted outlier trials. Weights were determine for each condition separately so as not to unduly distort signal in unexpected trials which, by definition, had fewer trials than the expected condition. Averages were then filtered with a low-pass filter with a cut-off of 30Hz to remove high frequencies introduced by the robust averaging method. These preprocessed signals were used in all of the analyses reported later on in the manuscript. They were only additionally manipulated in the SPM analysis, as reported below.

#### Analysis of habitation

We obtained ERPs time-locked to taste delivery to test for ERP habituation to the actual delivery of taste over time. For the purpose of the habituation analysis we analysed response to the taste itself (rather than to the cue). Although there is a risk that signal-to-noise ratio may be reduced in taste ERP trials due to muscle artefacts associated with the stimulus, previous research has confirmed an ERP response to taste at electrodes Fz, F3, F4, Cz, C3 and C4 (Franken et al., 2011; Kobal, 1985). We averaged across these electrodes within the window of taste delivery (0-1000ms after taste onset) for the first and second half of the experiments for the four conditions where taste was delivered, regardless of how expected the taste was. These data were analysed in SPSS entered into a 2 (taste) x 2 (experiment half) repeated-measures ANOVA using a threshold of p < .05.

#### Difference wave analysis of the FRN

Firstly, we conducted a difference-wave analysis to facilitate comparison with previous results. We extracted data from vertex electrodes Fz, FCz and Cz, 240-340ms after the outcome cue, following the recommendations of a meta-analysis which identified this spatiotemporal window as the most likely latency and location of the FRN signal of prediction error (Sambrook & Goslin, 2015a). Difference waves representing unexpected outcomes were computed using the conventional loss-minus-gain technique. In sweet we computed the omission-delivery difference wave, and in bitter we computed the delivery-omission difference wave. We conducted a one-sample t-test on this signal, following the analysis protocol of a recent study of the FRN to appetitive and aversive outcomes using a threshold of p < .05 (Heydari & Holroyd, 2016).

#### SPM analysis of frontocentral electrodes in the 200-380ms time window

The FRN literature lacks consistency in measuring the FRN. The meta-analysis we relied on to compute the difference waves above (Sambrook & Goslin, 2015a) acknowledged that variability in methods may complicate a blanket application of that time window. Though the FRN is linked to activity in the ACC (Walsh & Anderson, 2012) and well-defined as being expressed at frontocentral electrodes after 200ms, the specific electrode and latency varies between studies, meaning the peak signal may be overlooked. This is a particular issue in more complex experimental designs or where novel modalities, such as pain, could change the morphology of the signal (Garofalo et al., 2014; Talmi et al., 2013). Because we used a novel feedback modality here (taste), and because the difference-wave approach does not allow a test of the predictions of the axiomatic model (Talmi, Fuentemilla, Litvak, Duzel, & Dolan, 2012), we employed an additional data-driven approach for data analysis.

Global Field Power (GFP) is a technique that measures variance across all electrodes, conditions, and participants, and has traditionally been used to select spatiotemporal analysis windows (Lehmann & Skrandies, 1980; Skrandies, 1990). Using GFP we observed two peaks in frontocentral ERP activity, where the FRN is normally expressed (Walsh & Anderson, 2012), 200-380ms from the outcome cue (figure 2). This window was used for statistical analysis with SPM, an established technique (Litvak et al., 2011) which employs the General Linear Model to estimate parameters over electrodes and time, and which has been successfully used to study the FRN (Hauser et al., 2014; Litvak et al., 2011; Talmi et al., 2013). For this analysis the preprocessed data were converted to a single three-dimensional space by time image was created for each subject and condition. This conversion is achieved by generating a scalp map for each condition and stacking these maps over peristimulus time. The resulting images were smoothed using a Gaussian kernel full-width at half maximum of 8 mm/ms. Individual smoothed images for each condition were entered into a two statistical models, one for each taste, and analysed with a 2 (expectancy: expected/unexpected) x 2 (outcome: delivered/omitted) repeated-measures ANOVA. Higher-order effects were analysed first, and, used to mask exclusively the analysis of lower-order effects. Following (Talmi et al., 2013), a peak threshold of p < 0.005 and a cluster extent threshold of 100 voxels was used. All key results are reported corrected for multiple comparisons at the cluster level using a strict FWE < .05.

**Figure 2:**
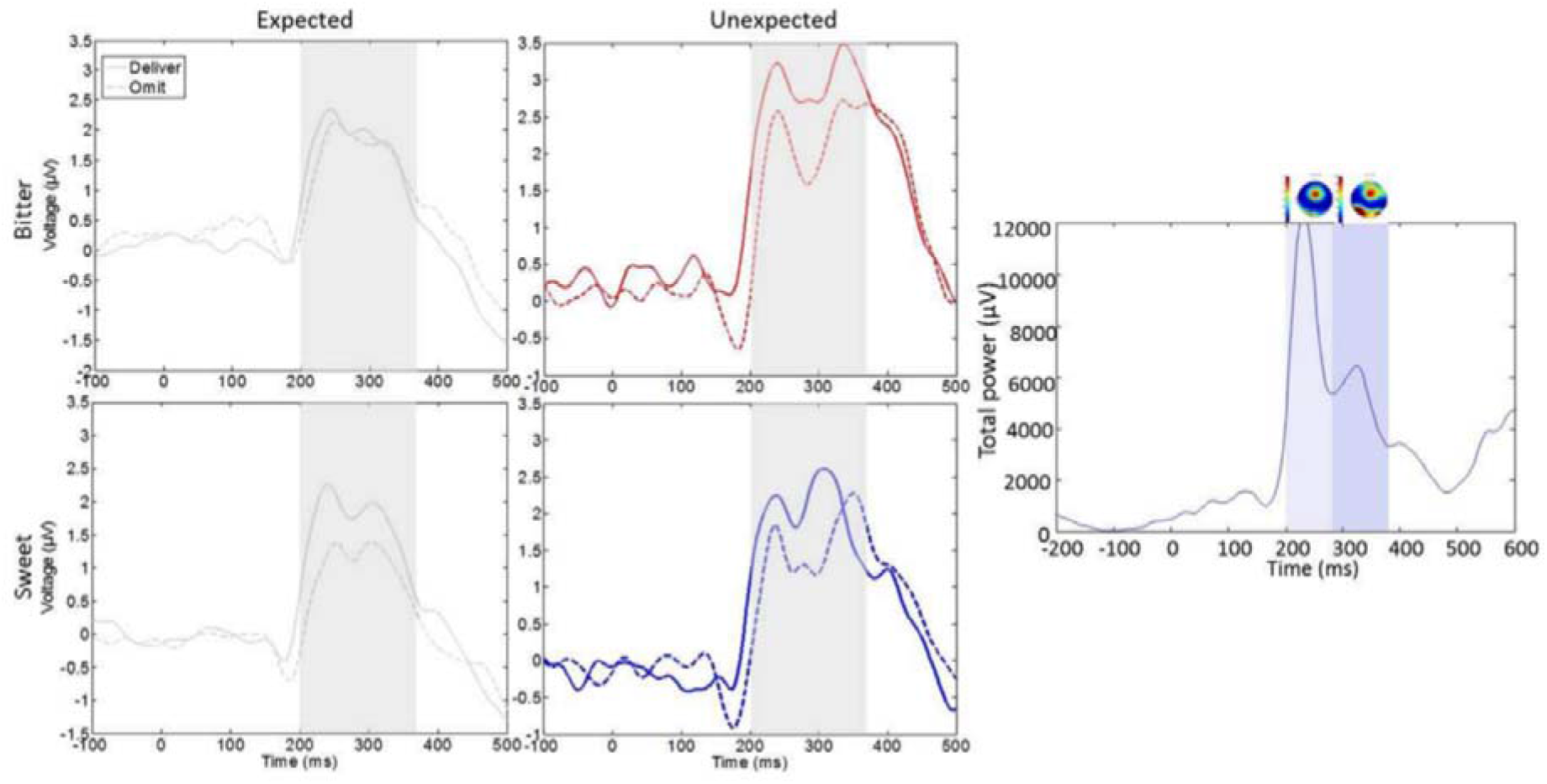
Detailed results. Top: group average ERPs for all conditions. ERPs were averaged over frontocentral electrodes and time-locked to the outcome cue presented at 0ms. *Bottom:* G*lobal field power*. Global field power identified two peaks in time windows 200 to 380ms post outcome cue. Topographic plots at the top of this panel show the topography of the global field power for the two time-windows.

## Results

### Behavioural results

We ran a 2 (magnitude: high, low) x 2 (taste: bitter, sweet) factorial ANOVA on the ratings of intensity for bitter and sweet for the concentrations we selected for each participant. Unsurprisingly, there was a significant main effect of magnitude on the ratings, where high magnitude tastes were rated as significantly more intense than low 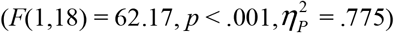. There was no significant effect of taste 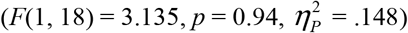, and no interaction 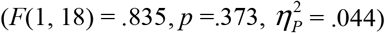. Participants were 97% accurate in identifying whether or not they had seen a cue in the previous 40 trials, which confirmed that they were attending to the cues.

### Electrophysiological results

#### Habituation

Taste-elicited amplitudes showed no significant effect of taste 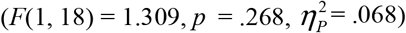, experiment half 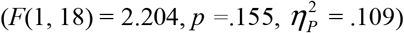, or interaction 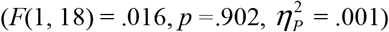, suggesting participants did not habituate to the taste.

#### Difference-wave analysis of the FRN

Figure 3 depicts a significant FRN in sweet (*t* (18) = −2.152, *p*=.045), replicating previous work. Crucially, in bitter we observed a significant ‘reverse’ FRN (*t* (18) = 2.78, *p* = .012), replicating our previous findings with pain (Talmi, Anderson & El-Deredy, 2013).

**Figure 3:**
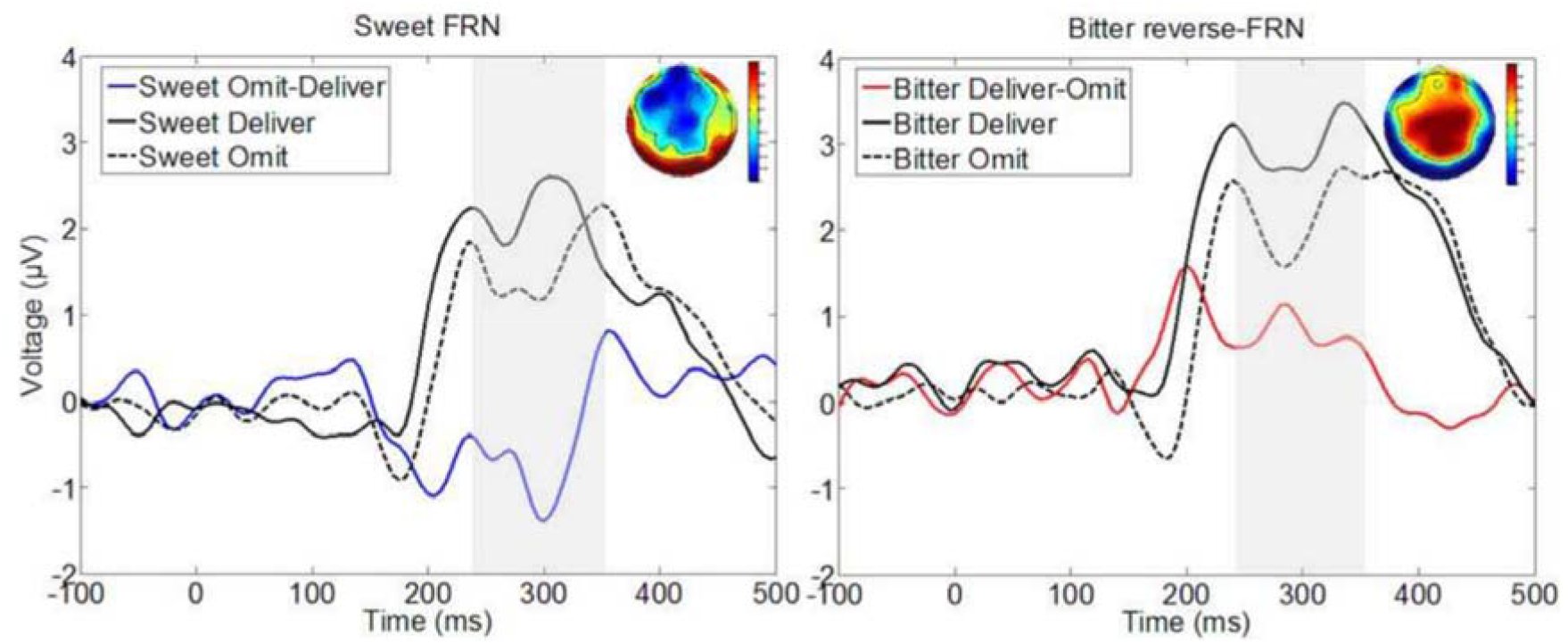
Difference-wave FRN analysis. Difference waves averaged over Fz, FCz and Cz across participants, subtracting unexpected loss from unexpected gain. The difference waves are time-locked to the outcome cue presented at 0ms. The search volume (240-340ms), based on a meta analysis, is shaded. Topographic plots show the unexpected loss-gain difference wave topographies for the shaded time-window.

### SPM analysis of frontocentral electrodes in the 200-380ms time window

#### Aversive and reward prediction error

The outcome by expectancy interaction was analysed for each taste. Figure 4 shows that the pattern of this interaction was similar in both tastes, but more pronounced in bitter. In bitter, the interaction was expressed in a significant cluster peaking at 285ms, which extended 251-317ms after the outcome cue (peak at C2, *x* = 17.00, *y* = −3.25, cluster *p* (FWE) = .032, cluster size 618 voxels). We conducted further analyses to unpack this interaction. Masked inclusively by the interaction of outcome and expectancy, unexpected delivered outcomes were significantly more positive than omitted (peak at C2, *x* = 12.75, *y* = −8.63, extending 200-368ms after the outcome cue, cluster *p* (FWE) <.001, cluster size 2855 voxels). There was no significant difference between expected delivered and omitted outcomes. Unexpected delivered outcomes were significantly more positive than expected (peak at Cz, *x* = 0.00, *y* = −8.63, extending 200-380ms after the outcome cue, cluster *p* (FWE) = .003), and unexpected omitted outcomes were significantly more negative than expected (peak at FC4, *x* = 34.00, *y* = 7.50, extending 269-295ms after the outcome cue, peak *p* (FWE) < .001). As shown in figure 4, this funnel-shaped signal adhered to the criteria of an axiomatic model of prediction error in the aversive domain. In Sweet the outcome by expectancy interaction did not survive our significance threshold. It can be readily appreciated that the numerical pattern of the nonsignificant interaction in sweet contradicts an interpretation of the frontocentral signal as a utility prediction error, because in both tastes unexpected delivered outcomes – good in sweet, bad in bitter-yield more positive amplitudes than unexpected omitted outcomes.

**Figure 4:**
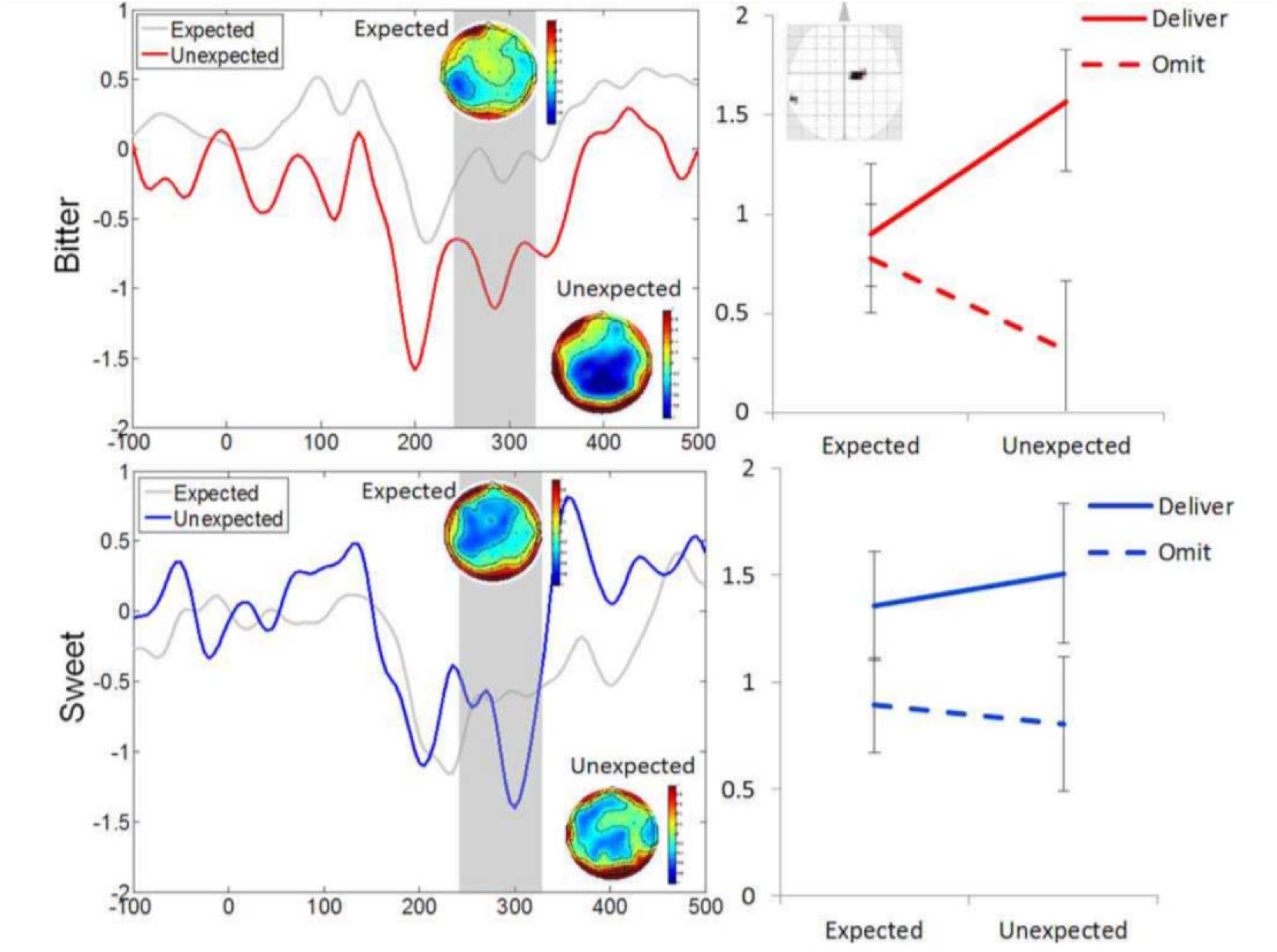
Aversive prediction error but no reward prediction error. Left: Difference waves subtracting delivered from omitted outcomes. The difference waves were averaged over frontocentral electrodes across participants, and time-locked to the outcome cue presented at 0ms. The temporal boundaries of the significant interaction are shaded in grey. In the Sweet condition (lower left) the interaction was not significant. Right: Voltages at the peak voxel of the cluster (inset, plotted on the SPM glass brain) corresponding to the significant Outcome-By-Expectancy interaction cluster in the Bitter condition (upper right) and the nonsignificant Outcome-By-Expectancy interaction in the Sweet condition (lower right). Error bars represent the standard error of the mean.

#### Motivational salience

The analysis of the main effect of outcome in each taste was masked exclusively by the interaction of outcome and expectancy. In bitter, the effect of outcome yielded a frontocentral cluster where amplitude for delivered outcomes was more positive than for omitted outcomes, peaking at 215ms and extending 200-287ms after the outcome cue (peak at FC1, *x* = −21.25, *y* = 2.13, cluster *p* (FWE) < .001, cluster size 3582 voxels). A similar result was obtained in sweet, where a frontocentral cluster peaked at 220ms and extended 200-265ms after the outcome cue (peak at C1, *x* = −17.00, *y* = −3.25, cluster *p* (FWE) = .001, cluster size 2181 voxels).

#### Surprise salience

The analysis of the main effect of expectancy in each taste was again masked exclusively by the interaction of outcome and expectancy. In bitter, the effect of expectancy was expressed in a frontocentral cluster peaking at 370ms and extending 306-380ms, where amplitude for unexpected outcomes was more positive than amplitude for expected outcomes (peak at FCz, *x* = −8.50, *y* = 7.50, cluster *p* (FWE) <.001, cluster size 2861 voxels). Again, a similar result was obtained in sweet, where the effect of expectancy yielded a frontocentral cluster peaking at 375ms (peak at C1, *x* = −8.5, *y* = −3.25, 309-380ms, cluster *p* (FWE) <.001, cluster size 2525 voxels).

#### Analysis across tastes

The analyses above yielded a number of similarities across tastes which are explored here with a more comprehensive model, including taste as an additional within-subject factor. Although the Outcome-by-expectancy interaction was of different magnitudes in bitter and sweet, the three-way interaction between outcome, expectancy, and taste was not significant. The two-way outcome-by-expectancy interaction (across bitter and sweet tastes) yielded a frontocentral cluster peaking at 285ms which extended 273-317ms after the outcome cue (peak at C2, *p* < .001, *x* = 21.25, *y* = −3.25). While significant, this activation did not survive our conservative correction for multiple comparisons; this is clearly because the weak interaction in sweet attenuated the highly significant interaction in bitter. The main effect of outcome, masked exclusively by the interaction of outcome and expectancy, yielded a frontocentral cluster peaking at 220ms and extending 200-342ms after the outcome cue (peak at C1, *x* = −21.25, *y* = −3.25, cluster *p* (FWE) < .001, cluster size 4168 voxels). The main effect of expectancy masked exclusively by the interaction of outcome and expectancy yielded a frontocentral cluster peaking at 370ms and extending 203-380ms after the outcome cue (peak at FCz, *x* = −4.25, *y* = 2.13, cluster *p* (FWE) < .001, cluster size 3902 voxels).

## Discussion

This is the first study to characterise ERP expression of prediction error using taste, thus bridging a gap between human EEG studies on reinforcement learning and those carried out in animal models (Holroyd & Coles, 2002; Schultz, Dayan, & Montague, 1997). We delivered and omitted expected and unexpected hedonically matched appetitive and aversive tastes. Our goal was to distinguish between utility prediction error signals, where amplitude should be most positive (or negative) for delivered appetitive and omitted aversive tastes and most negative (or positive) for omitted appetitive and delivered aversive tastes, and a salience response, where amplitude should be most positive (or negative) for the delivery of both appetitive and aversive tastes compared to their omission. Following the logic of the axiomatic model (Caplin & Dean, 2008; Rutledge et al., 2010) we were particularly interested in signal that differentiated aversive and appetitive taste and taste omission cues more strongly when they were unexpected. For both appetitive and aversive taste, we observed continued expression of salience (Sambrook & Goslin, 2015b) across the entire 200-380ms time window. There was also an expression of aversive but not reward prediction error at the latency most characteristic of the FRN, peaking at 285ms (Holroyd, 2004; Sambrook & Goslin, 2015a).

The latency of the FRN, considered the ERP correlate of prediction error, is variable between studies and is usually identified between 200-350ms after the outcome (Hauser et al., 2014; Heydari & Holroyd, 2016; Holroyd, Hajcak, & Larsen, 2006; Holroyd, 2004). The lack of consistency in selection of latency windows makes it difficult to conclude whether the same signal is being studied across laboratories. A meta-analysis suggested that frontocentral vertex electrodes are most likely to express reward prediction error, mainly related to monetary reward, around 240-340ms, although the latency could be influenced by the modality of the outcome and other features of the experimental set-up (Sambrook & Goslin, 2015a). Here we examined the ERPs from 200ms onwards, firstly employing the traditional difference-wave approach within the electrodes and time-window recommended in the meta-analysis, and secondly in through a data-driven analysis in the spatiotemporal window of 200-380ms identified through coarse GFP analysis.

The commonly-used loss-gain difference-wave analysis (Heydari & Holroyd, 2016) was conducted to facilitate comparison between this study and previous FRN studies. We observed significant expression of appetitive FRN and aversive reverse-FRN. This result was driven by an increased positivity in response to delivered over omitted tastes in both appetitive and aversive domains and directly replicates our previous work, where we observed increased positivity to the delivery of pain and money (Talmi et al., 2013), with a reverse-FRN for pain. The implication of these results is that the FRN may express response to more and less salient stimuli, rather than differentiate ‘bad’ from ‘good’.

In the data-driven analysis ERPs had a more positive amplitude for tastes that were about to be delivered compared to those that were about to be omitted, and were also more positive for tastes that were unexpected compared to those that were expected. Using the terminology of Schultz (2016), this expression of motivational salience and surprise salience, respectively, was observed for both appetitive and aversive outcomes (i.e. sweet and bitter tastes). The response to motivational salience peaked particularly early, around 220ms, in agreement with recent proposals of an initial unsigned response that represents various forms of salience rather than utility (Schultz, 2016). This finding replicates our previous work with pain and money outcomes, where ERPs differentiated between pain and money that were about to be delivered and those about to be omitted around 200-290ms after the cue (Talmi et al., 2013). It is important to note that the main effect of outcome here drove the results reported above for the traditional difference-wave analysis. As per previous suggestions (Holroyd, 2004; Sambrook & Goslin, 2015b) the early response to motivational salience is likely related to the N2.

Frontocentral ERPs at 285ms in the aversive domain, in experimental blocks where only Delivery or Omission of bitter taste was possible, conformed to the logic of an axiomatic model of prediction error. The key pattern specified by the Axiomatic Model of prediction error is an interaction between outcome and expectancy (Rutledge et al., 2010). At that time window in the bitter condition ERPs were more positive for outcome delivery relative to omission, a difference that was more pronounced when outcomes were unexpected. The signal was also significantly more positive for delivered unexpected than delivered expected outcomes, and more negative for omitted unexpected than expected outcomes, in line with the predictions of the axiomatic model. On its own, as discussed earlier, the signal observed in the bitter condition adheres to criteria for an aversive prediction error, but cannot be interpreted unambiguously as a utility prediction error signal.

In the appetitive domain at that same latency, in blocks where only delivery or omission of sweet taste was possible, the signal followed a similar pattern to that found in the aversive domain, but while the main effects of outcome and expectancy were significant, the interaction between them was not. Two aspects of this result are important. First, because this outcome by expectancy interaction is a hallmark of a neurobiological reward prediction error signal we conclude that we could only observe an aversive prediction error here, but not a reward prediction error. Clearly, the finding that frontocentral ERPs did not express prediction error in the appetitive domain already means that they also did not express utility prediction error, a quantity which should be equally well observed in both appetitive and aversive domain; yet the null interaction between outcome and expectancy in sweet could be just due to low power. Crucially, a utility prediction error signal should present with an *opposite* sign in each domain, but the direction of averages in all four conditions was the same. Clearly, a neurobiological signal of utility prediction error would not be expressed as greater positivity for both unexpectedly good and bad outcomes. Taken together, the frontocentral signal observed in this experiment did not express a utility prediction error. Other aspects of the data suggest that ERPs also did not track utility *per se*. For example, ERPs did not differentiate expected omission and delivery of a bitter taste (which have different utility), but did distinguish between expected and unexpected sweet taste (which have the same utility).

We did not observe a reward prediction error pattern in the appetitive domain, although such a pattern has been readily observed in previous work (Sambrook & Goslin, 2015a; Walsh & Anderson, 2012) including in our own previous study (Talmi et al., 2013). Instead ERPs in the appetitive domain expressed motivational and surprise salience. This may be due to the reinforcement modality we selected. Very few ERP studies of FRN used primary reinforcers. Although some previous ERP studies of aversive prediction error used pain, which is a primary reinforcer (Garofalo et al., 2014; Heydari & Holroyd, 2016; Talmi et al., 2013), all previous ERP studies of reward prediction error, including those that used pain in the aversive domain, used money, a secondary reinforcer, in the appetitive domain. We used primary taste reinforcers because appetitive taste is known to elicit a dopamine response. Second-by-second dopamine release in response to food cues signals future appetitive outcomes (Hamid et al., 2015), and anticipation of appetitive taste activates the dopaminergic system (O’Doherty, Deichmann, Critchley, & Dolan, 2002). In agreement with previous research (Nitschke et al., 2006), we did not see neural habituation to appetitive taste. Moreover, we followed routine practice in animal models and ensured that the hedonic value of our taste reinforcers was titrated so that it was positive and high for each individual participant, and we deprived participants of food and water beforehand, which enhanced the incentive salience (Berridge, 2012; McClure, Daw, & Montague, 2003) of the sweet taste. We propose, therefore, that the sweet taste is more salient than the small amounts of money participants gained in previous work, and so ERPs to sweet taste prioritised expression of salience, rather than prediction error. This hypothesis can be tested in future work where monetary and taste reinforcers are directly compared. We also acknowledge that it is possible, as the use of sweet taste as an appetitive reinforcer in humans is fairly novel, that the sweet taste was salient, but not appetitive, despite titration of the hedonic intensity of the sweet taste, and so elicited a salience response. However, previous research has successfully used sweet appetitive taste reinforcers (Franken et al., 2011; Kim, Shimojo, & O’Doherty, 2011; McClure, Berns, & Montague, 2003), so this hypothesis remains questionable. Future studies could acquire trial-by-trial ratings of hedonic intensity and appetitive value of sweet taste to test this.

We employed a Pavlovian task because this task established the fundamental reward prediction error result in animal models, which we aimed to be closest to. However, the fact that we employed a passive Pavlovian task, in contrast to the majority of FRN experiments which tend to employ instrumental learning tasks, could also contribute to the lack of reward prediction error signal. This is in keeping with previous work showing a smaller effect of reward prediction error in passive tasks (Sambrook & Goslin, 2015a). Therefore, it is possible that the FRN signals an instrumental, rather than a Pavlovian, reward prediction error. The fact that the ERPs signalled prediction errors in the aversive domain clarifies that domain cannot be ignored in the pursuit of an ERP signature of prediction error.

As we did not use source localisation analysis, we cannot make claims about where the signal originated from. A number of different regions expressed salience in previous fMRI work, summarised in a recent meta-analysis (Lindquist et al., 2015). The most likely sources of the salience response are the ACC, which receives prediction error signals from the midbrain, and the insula, which together with the ACC form the salience network (Seeley et al., 2007). The ACC was the source of the frontocentral salience response to money in a previous ERP-fMRI study, and this signal was shown to be signalled directly from dopaminergic sources to the ACC (Hauser et al., 2014). Furthermore, the ACC is known to be the source of the FRN ERP (Miltner, Braun, & Coles, 1997; Walsh & Anderson, 2012). The anterior insula and striatum have also been shown to express salience to appetitive and aversive tastes (Metereau & Dreher, 2013). The aversive prediction error signal we recorded on the scalp may have originated from the dopaminergic midbrain, involved in previous studies of aversive prediction error (Brischoux et al., 2009; Seymour et al., 2004), but its pathway to influencing scalp ERPs awaits further work.

Our findings go beyond existing fMRI work to exploit the temporal resolution of EEG and contradicts the dominant perspective on the FRN ERP as a signal of reward prediction error (Abler, Walter, Erk, Kammerer, & Spitzer, 2006; Carlson, Foti, Mujica-Parodi, Harmon-Jones, & Hajcak, 2011; Hauser et al., 2014; Holroyd, 2004; Holroyd & Coles, 2002; Walsh & Anderson, 2012). They agree better with recent work that proposes this signal expresses salience (Garofalo et al., 2014; Hauser et al., 2014; Huang & Yu, 2014; Pfabigan et al., 2015; Sambrook & Goslin, 2015b; Talmi et al., 2013). Our findings also agree with the PRO model, which asserts that ACC activity reflects negative surprise, responding to the unexpected omission of both appetitive and aversive outcomes (Alexander & Brown, 2011), in that ERPs were more negative to omissions across the board.

Our data show conclusively that frontocentral ERPs at the time-window of the FRN does not express outcome valence, contradicting the interpretation of the FRN as a utility prediction error signal (Holroyd, 2004). Across the time window of interest and across tastes ERPs were more positive for cues that predicted salient outcomes, namely, delivered or unexpected outcomes. The spatiotemporal evolution of the signal was differentially sensitive to the feature that rendered the predicted outcome salient, with a particular time window where the response in the aversive domain appeared to go beyond salience to resemble an aversive prediction error.

## Author notes

## Acknowledgements

This work was supported by a studentship grant from the Medical Research Council, UK. WeD acknowledges the support of CONICYT, Chile, Basal project FB0008 and FONDECYT project 1161378. We thank O. Adams for her assistance with data collection. WeD is currently affiliated with the School of Biomedical Engineering, University of Valparaiso, Chile. We declare no conflicts of interests.

## Name and address for reprints

Miss Emily Hird, Zochonis building, Division of Neuroscience and Experimental Psychology, University of Manchester, United Kingdom, M139GB.

